# Seizure control via astrocyte targeted optogenetic extracellular potassium modulation: computational analysis of K^+^ vs non-selective cation conducting opsins

**DOI:** 10.64898/2026.09.24.754075

**Authors:** Laila Weyn, Thomas Tarnaud, Wout Joseph, Robrecht Raedt, Emmeric Tanghe

## Abstract

**Aims:** Seizure generation has been strongly linked to the accumulation of extracellular potassium. As astrocytes play a key role in potassium homeostasis, astrocyte targeted optogenetic modulation of extracellular potassium has been proposed as a novel strategy for seizure control. The effectiveness of such modulation is expected to depend strongly on the ion selectivity of the employed opsin. This study aims to mechanistically compare potassium-selective (WiChR) and cation-conducting (ChR2-like, ChR2^*∗*^) opsins for astrocyte targeted modulation of extracellular potassium, and to determine how astrocytic properties and stimulation parameters influence modulation efficacy.

**Methods:** A conductance-based computational framework was used to model astrocyte extracellular space interactions, incorporating ion-specific photocurrents, Na^+^*/*K^+^-ATPase activity, and gap junction-mediated coupling within the astrocytic syncytium. Systematic parameter sweeps were performed to evaluate the influence of ion selectivity, astrocytic parameters, and stimulation paradigms on potassium dynamics.

**Results:** WiChR-based modulation consistently produced more effective and robust reductions in [K^+^]_e_ than ChR2^*^, via direct K^+^ inward currents and reduced dependence on pump activity. In contrast, ChR2^*^ primarily limited potassium accumulation rather than reversing it and exhibited greater sensitivity to parameter variations, particularly Na^+^*/*K^+^-ATPase activity. The effectiveness of astrocyte targeted modulation was strongly dependent on syncytial integrity, with reduced gap junction coupling impairing potassium clearance. Optimal stimulation performance for WiChR was observed for a pulse repetition period of 20 *ms*, with outcomes determined by the interaction between opsin kinetics and stimulation parameters.

**Conclusions:** These results demonstrate that activation of WiChR opsins in astrocytes can reduce extracellular potassium levels and thereby modulate neuronal excitability. Ion selectivity emerges as a key determinant of modulation efficacy at the astrocytic level, with potassium-selective opsins providing a more direct and reliable approach than non-selective cation-conducting alternatives. Together, these findings establish a mechanistic basis for optogenetic activation of astrocytic potassium buffering as a potential strategy for seizure suppression and other neurological disorders associated with potassium dysregulation.

**AUTHOR SUMMARY:** Excessive accumulation of potassium in the extracellular space is thought to contribute to seizures by increasing neuronal excitability. Astrocytes play a central role in regulating extracellular potassium levels and have therefore emerged as promising targets for therapeutic intervention. Optogenetic tools enable light-controlled ion currents to be introduced into astrocytes, potentially enabling direct modulation of potassium homeostasis. Previous experimental work has shown that astrocytic modulation via non-selective cation channels can reduce seizure activity, but the recent development of potassium-selective opsins raises the possibility of directly controlling astrocyte-mediated potassium currents. However, it remains unclear how the ion selectivity of these currents influences their ability to regulate extracellular potassium.

Using a computational model of astrocyte potassium homeostasis, we compared the effects of activating a potassium-selective opsin (WiChR) with a non-selective cation-conducting opsin (ChR2-like). We found that potassium-selective modulation produces a larger and more robust reduction in extracellular potassium because potassium currents directly contribute to potassium clearance. In contrast, ChR2-like modulation relies more strongly on indirect mechanisms, including sodium accumulation and enhanced pump activity, making its effectiveness more sensitive to cellular properties and network conditions. We also identified astrocytic gap-junction coupling as a key determinant of successful potassium regulation.

These findings identify potassium-selective opsins as a promising tool for controlling extracellular potassium dynamics. More broadly, this study provides mechanistic insight into how astrocytes regulate brain excitability and offers guidance for the development of future optogenetic strategies targeting disorders associated with disrupted potassium homeostasis.

## Introduction

Temporal lobe epilepsy (TLE) is a chronic neurological disorder characterized by recurrent seizures originating in the temporal lobe. Although anti-seizure medication remains the primary treatment, a substantial proportion of patients are drug-resistant, highlighting the need for alternative therapeutic strategies^1–3^. Electrical neuromodulation approaches such as deep brain stimulation (DBS) and responsive neurostimulation (RNS) have demonstrated clinical efficacy, but their limited cell type specificity can result in the unintended activation of off-target neuronal populations and fiber tracts that are not directly involved in seizure generation^2,4,5^.

Optogenetics offers a promising alternative by enabling cell type specific and spatially precise modulation of neural activity using light-gated ion channels^6–8^. Depending on their ion selectivity, opsins can induce neuronal excitation or inhibition upon illumination. While early optogenetic seizure suppression strategies primarily focused on neuronal populations, either through direct inhibition of excitatory neurons or activation of inhibitory interneurons, astrocytes have increasingly emerged as attractive targets for optogenetic intervention and other novel therapeutic strategies^9,10^. Astrocytes play a central role in maintaining extracellular potassium (K^+^) homeostasis, which is critically implicated in seizure dynamics^10–14^. Indeed, extracellular potassium concentration ([K^+^]_e_) has been proposed as a key ictogenic factor due to its strong influence on neuronal excitability and experimental evidence linking [K^+^]_e_ dysregulation to seizure initiation and propagation^11–15^.

Astrocytes regulate extracellular K^+^ through multiple mechanisms, including inward-rectifying potassium channels (Kir4.1), Na^+^*/*K^+^-ATPase activity, and redistribution via gap junction-mediated spatial buffering within the astrocytic syncytium, the network of interconnected astrocytes that enables isopotentiality^16–19^. Building on this physiological role, Zhao et al. (2022) demonstrated that optogenetic stimulation of Channelrhodopsin-2 (ChR2) expressing astrocytes can reduce seizure activity by increasing intracellular Na^+^, thereby enhancing Na^+^*/*K^+^-ATPase function and promoting clearance of K^+10^. However, this approach relies on an indirect mechanism in which Na^+^ influx modulates potassium handling via metabolic and pump-dependent processes, potentially introducing confounding effects such as altered membrane potential, ionic imbalance, or increased energetic demand^10^.

Potassium-selective opsins, such as Wobblia inhibitory channelrhodopsin (WiChR), offer a fundamentally different strategy by directly mediating K^+^ flux across the membrane^20,21^. Unlike non-selective cation channels, which conduct a mixture of ions and whose photocurrents are typically dominated by Na^+^ influx because |*V*_*m,A*_ − *E*_*Na*_| ≫ |*V*_*m,A*_ − *E*_*K*_|, WiChR generates photocurrents carried predominantly by K^+^ ions. As a result, optogenetic activation of WiChR can directly augment astrocytic K^+^ buffering. However, under resting conditions the astrocytic membrane potential is typically slightly depolarized with respect to the potassium reversal potential, raising the possibility that activation could drive outward K^+^ currents and increase [K^+^]_e_^18^. Although under increased [K^+^]_e_ conditions, E_K_ is expected to increase and the astrocytes tend to maintain a stable membrane potential through gap-junction-mediated syncytial isopotentiality^17^. The resulting difference between *E*_*K*_ and *V*_*m,A*_ would generate an inward photocurrent upon optogenetic activation. This requires further investigation as it raises the question of whether direct potassium modulation provides advantages over indirect sodium-driven approaches.

Despite the growing interest in astrocyte targeted optogenetics, a direct comparison between cation-conducting (ChR2-like) and potassium-selective (WiChR) opsins in this context is currently lacking. In particular, the interplay between ion selectivity, astrocytic physiology, and extracellular potassium dynamics is complex and difficult to isolate experimentally, as these processes depend on tightly coupled ionic gradients, transport mechanisms, and network interactions that are challenging to measure *in vivo*. Computational modelling provides a powerful framework to address this challenge by enabling controlled manipulation of opsin properties, cellular parameters, and stimulation paradigms. Previous modelling studies have explored astrocytic contributions to potassium buffering and neuronal excitability^16,17,22^. These models can be combined with mathematical descriptions of opsin kinetics, initially developed for ChR2^23^ and later extended to study opsin dynamics in conductance-based neuron models^24,25^. The recently introduced opsin formalism by Weyn et al. (2025) enables direct parametrization from experimental photocurrent recordings, facilitating consistent comparison between different opsins^21^. Importantly, this framework can be extended to explicitly incorporate ion selectivity, thereby enabling mechanistic investigation of how permeability differences influence ionic dynamics^26^.

In this study, we use computational modelling to systematically compare astrocyte mediated optogenetic modulation using cation-conducting (ChR2-like) and potassium-selective (WiChR) opsins. We specifically investigate how ion selectivity shapes the capacity to modulate extracellular K^+^ and examine how astrocytic properties, opsin dynamics, and stimulation paradigms interact to determine the efficacy and directionality of potassium regulation.

## Methods and models

### Opsin model

The photocurrent generated by opsin channels is described using a variant of the double two-state opsin model (22OM), originally introduced by Schoeters et al. (2021) and modified by Weyn et al. (2025) (22OMs)^21,24^. In this study, we consider two opsin types with distinct ion selectivity properties: the potassium-selective opsin WiChR and a ChR2-like non-selective cation channel (denoted ChR2^*^ because WiChR kinetics are used to isolate the effect of ion selectivity). Both are modeled within the same formalism to enable a direct comparison of their effects.

In this framework, illumination at light intensity I triggers a transition of the channel from its closed to open configuration (C→O). Concurrently, light-dependent adaptation modifies the channel conductance, captured through the transition between the dark-adapted and light-adapted states (R→S). Once the light stimulus ends, the channel returns to the closed, dark-adapted state via the transitions O→C and S→R. The temporal evolution of these states is governed by the following differential equations.

In darkness (I = 0):

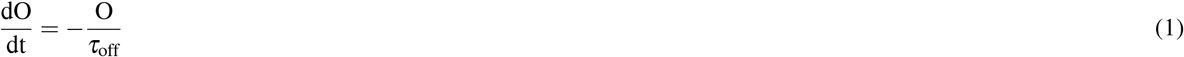

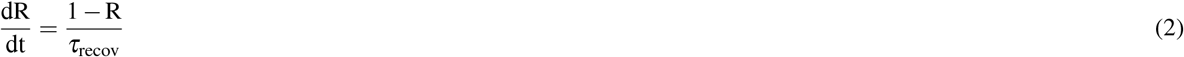

During illumination (I ≠ 0):

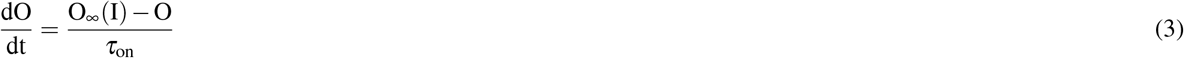

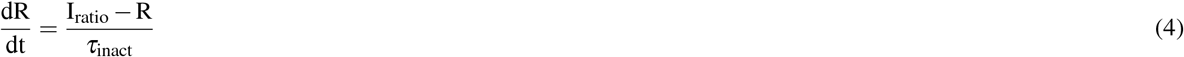

Here, O represents the fraction of channels in the open state (C=1-O), whereas R reflects the degree of dark adaptation (S=1-R), taking the value 1 under fully dark-adapted conditions and decreasing during illumination to represent reduced conductance in the light-adapted state. The photocurrent produced by an opsin type x is expressed as

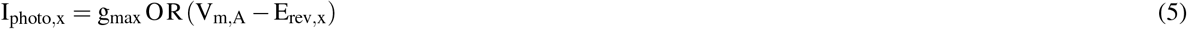

where g_max_ represents the maximal specific conductance in the fully open, dark-adapted state and E_rev,x_ denotes the opsin’s reversal potential. The rectification function G(V) from the original formulation (^24^) is omitted here, as the peak and steady-state currents exhibit an approximately linear dependence on the membrane potential, and thus (V − E_rev,x_) adequately captures voltage sensitivity. Model parameters can be inferred directly from experimental photocurrent recordings using the following measurable quantities and are reported in table 1^24^:

**Table 1.**
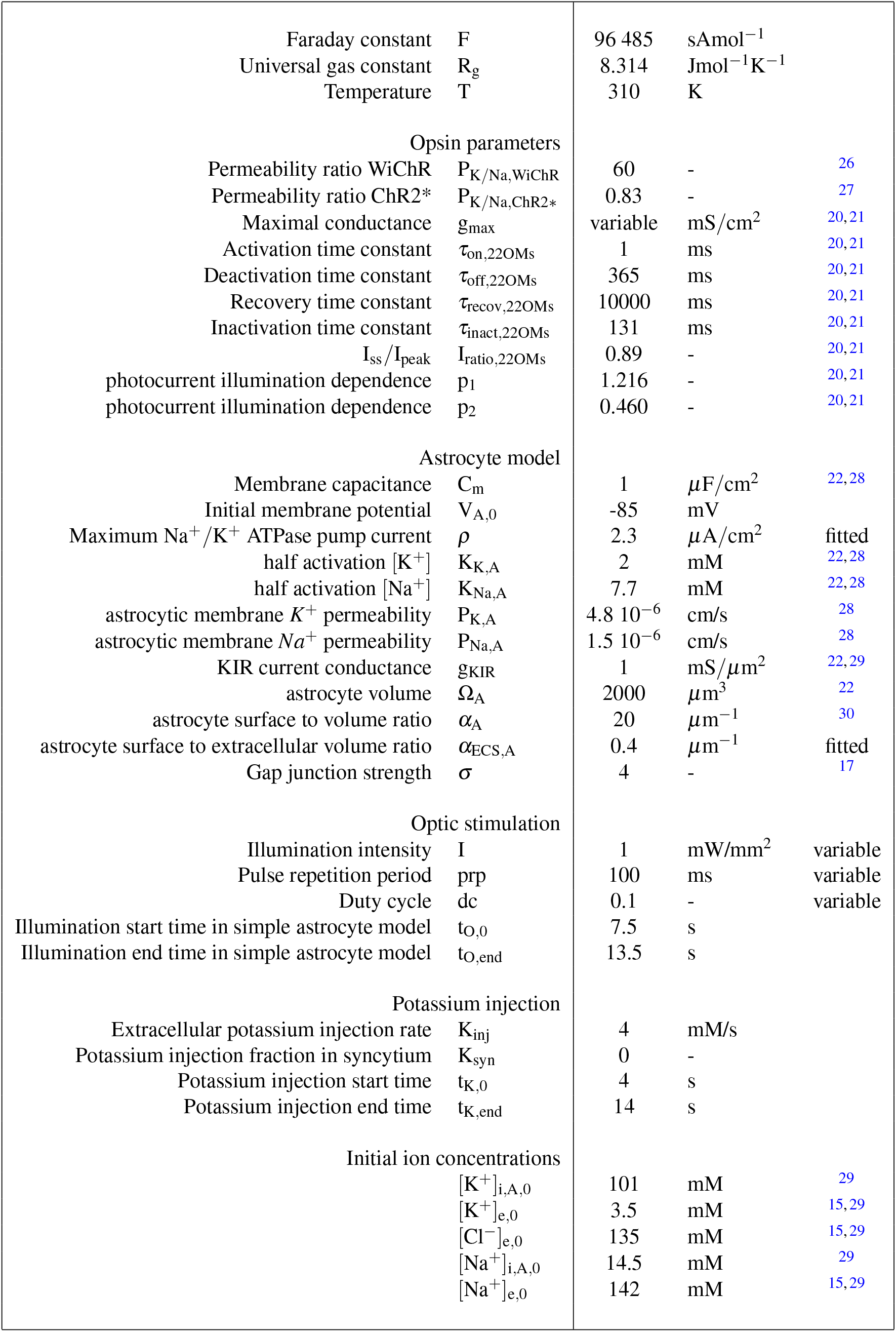
List of model parameters.

- Peak current (I_peak_): the maximal current observed during illumination.
- Steady-state current (I_ss_): the current reached in the light-adapted state; the ratio I_ratio_ = I_ss_*/*I_peak_ quantifies adaptation.
- Activation time constant (*τ*_on_): obtained from a mono-exponential fit of the rising phase.
- Deactivation time constant (*τ*_off_): obtained from a mono-exponential fit of the decay following light offset.
- Inactivation time constant (*τ*_inact_): obtained from a mono-exponential fit of the decay from I_peak_ to I_ss_.
- Recovery time constant (*τ*_recov_): obtained from mono-exponential fit of recovery of I_peak_ across repeated stimuli.
- Parameters p_1_ and p_2_ fitted to the intensity dependence:

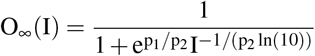

Parameter values were obtained by fitting the model to experimental photocurrent recordings of WiChR, while the same kinetic parameters were retained for ChR2^*^ to isolate the effect of ion selectivity^20,21^. The reversal potential E_rev,x_ is determined assuming conduction of both K^+^ and Na^+^ with permeability ratio P_K*/*Na,x_:

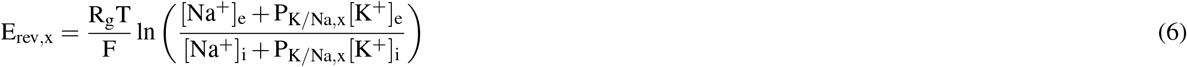

The key distinction between WiChR and ChR2^*^ is encoded in this permeability ratio, with P_K*/*Na,WiChR_ = 60 and 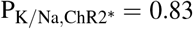 ^26,27^. Following Ohnemus et al. (2026)^26^, the total photocurrent is decomposed into K^+^ and Na^+^ components:

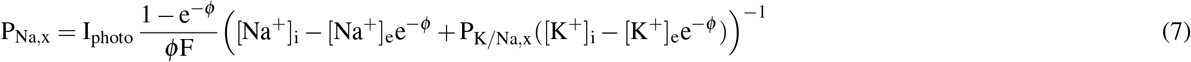

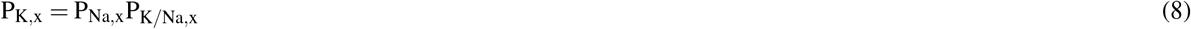

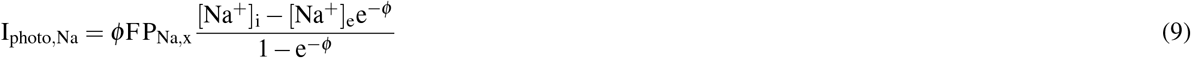

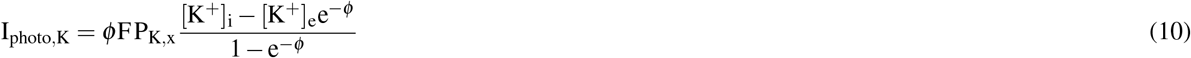

with 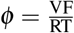. The fitted model is shown in Figure 1C.

**Figure 1.**
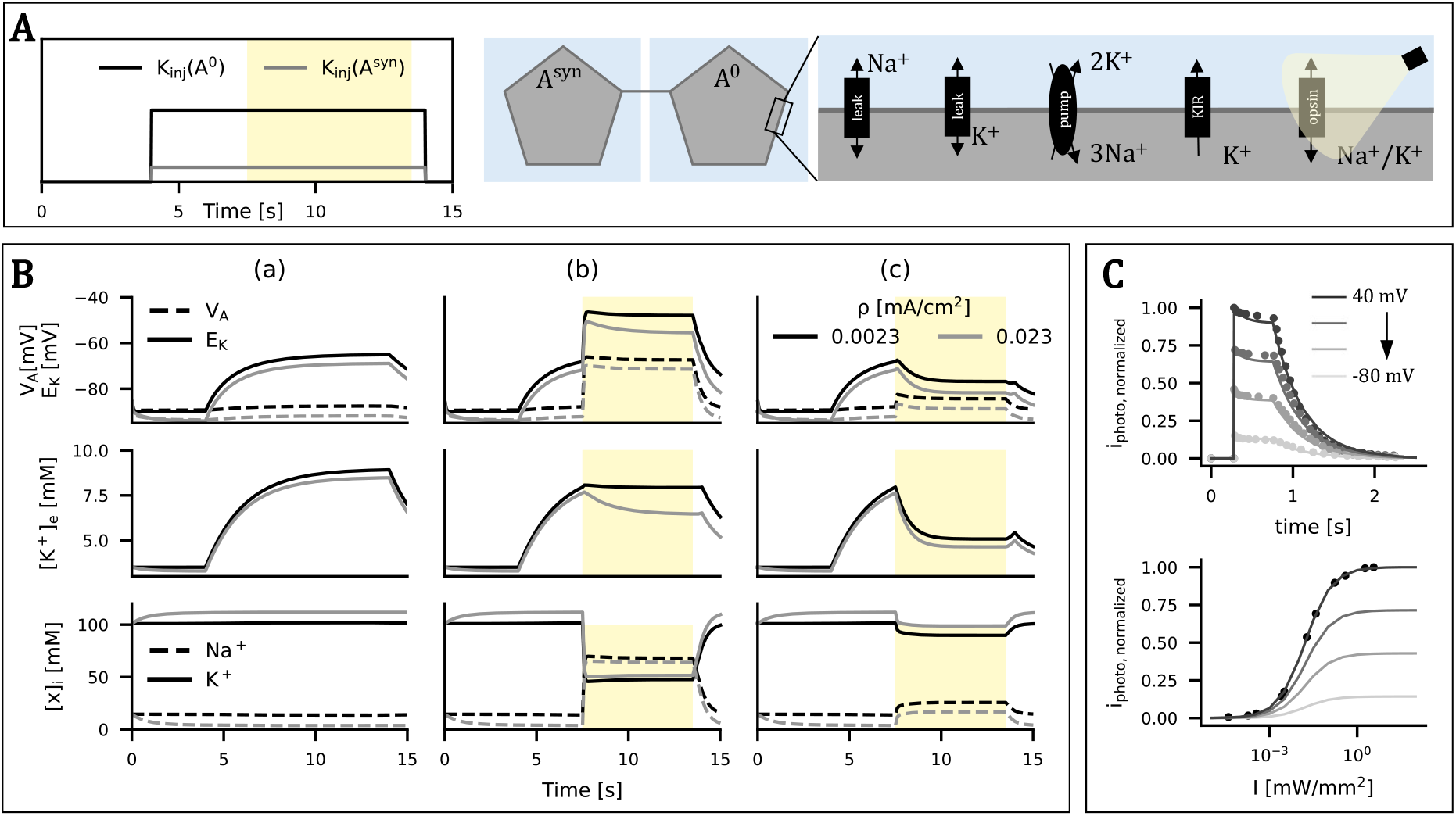
Model overview and stability. **A**. (left) Schematic timeline of potassium injection and optical stimulation. (right) Schematic representation of the gap-junction-coupled 2A model with all modelled currents. **B**. Output of the two-astrocyte model without optogenetic modulation (a) and with ChR2^*^ (b) or WiChR (c)-based modulation. **C**. Fit of the opsin model to experimental WiChR photocurrent data voltage, default I = 3.7 mW*/*mm^2^.

### Astrocyte model

To investigate how optogenetic modulation of astrocytes influences extracellular potassium dynamics, we employ a biophysically detailed astrocyte model. This model is based primarily on the single-compartment Hodgkin-Huxley type models introduced by Huguet et al. (2016) and Terman and Zhou (2019) for describing astrocytic membrane dynamics within neuron-glia networks^22,28^. In this formulation, the astrocyte membrane potential V_m,A_ evolves according to the combined contributions of potassium and sodium leak currents (I_K,A_, I_Na,A_), the Na^+^*/*K^+^ ATPase pump current (I_P,A_), an inward-rectifying potassium current (I_KIR,A_), and gap junction-mediated ionic exchange (I_gap_)^22^. The governing equation for the astrocyte membrane voltage is

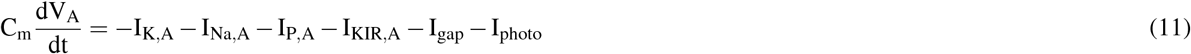

where C_m_ denotes the membrane capacitance. The potassium and sodium leak currents (I_K,A_, I_Na,A_) are expressed using the Goldman-Hodgkin-Katz (GHK) flux formalism taken from Huguet et al. (2016), yielding

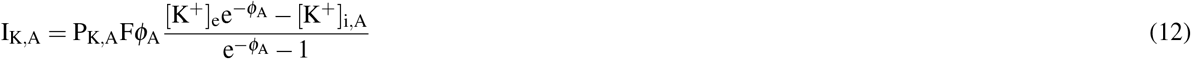

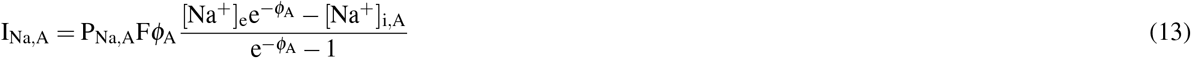

Here 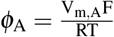, and P_K_ and P_Na_ represent the respective permeability coefficients. All parameter values are summarized in Table 1. The inward-rectifying potassium current I_KIR,A_ is defined as^22,31^:

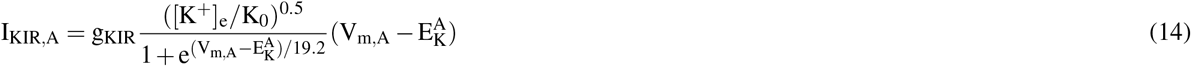

with K_0_ = 1 mM and 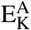 the *K*^+^ reversal potential. The Na^+^*/*K^+^ ATPase pump current retains the form used in both Terman and Zhou (2019) and Huguet et al. (2016):

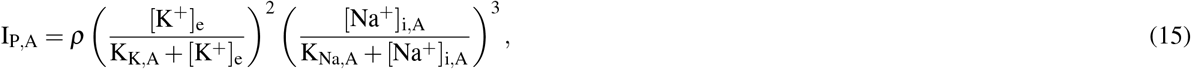

where *ρ* is the maximal pump strength, and K_K,A_ and K_Na,A_ are the half-activation constants. Gap junction coupling is modeled using the GHK formulation^17,28^. As illustrated in figure 1A, an astrocyte A^0^ is coupled to a representative syncytium astrocyte A^syn^, which does not receive any gap junction current and thus serves as a boundary condition representing the larger astrocytic network that maintains ion homeostasis. The K^+^ and Na^+^ components of the gap junction current are given by

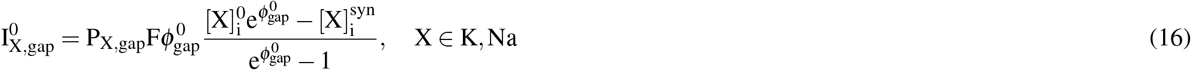

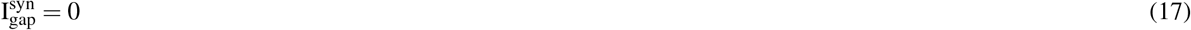

with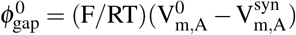. The corresponding permeabilities are^22,28^

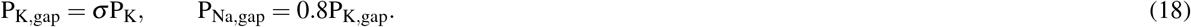

Here, *σ* indicates the gap junction strength.

Intracellular ion concentrations evolve according to conservation equations that incorporate membrane ionic fluxes and pump currents. Specifically,

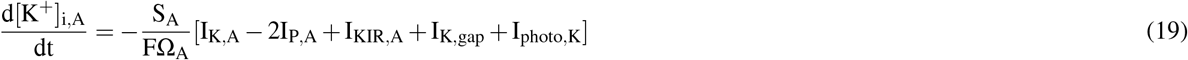

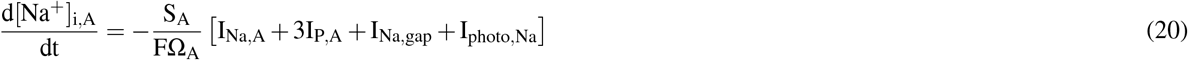

where Ω_A_ and S_A_ = *α*_A_Ω_A_ denote the astrocyte cytosolic volume and surface. The extracellular ion concentrations follow the following equations^22^

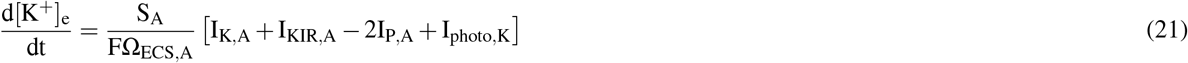

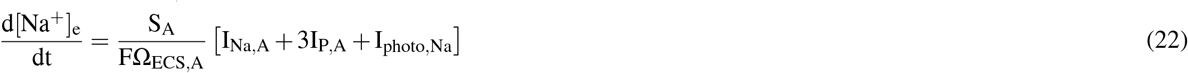

which reflect the combined influence of membrane fluxes and pump-mediated ion exchange on the extracellular space (ECS) compartment with volume Ω_ECS,A_ = S_A_*/α*_ECS,A_.

### Simulation setup

The modelled photocurrent was incorporated into the astrocyte model (A^0^, not A^syn^) to investigate how optogenetic modulation of astrocytes affects extracellular potassium concentration. To this end, a simulation setup was designed consisting of two astrocytes, A^0^ and A^syn^, each coupled to a separate ECS compartment (Figure 1A). [K^+^]_e_ in 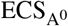 is increased at a constant rate K_inj_, while [K^+^]_e_ in 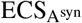 is increased at a rate of K_syn_.K_inj_. By default, K_syn_ = 0, such that the syncytium is unaffected. This increase either occurs between t_K,0_ and t_K,end_ (Ksteady = default setup), or between t_O,0_ and t_K,end_ with an instantaneous jump of [K^+^]_e_ to 8 mM at t_O,0_ (Kjump). Illumination for optogenetic modulation is applied between t_O,0_ and t_O,end_ (intervention) or throughout the entire simulation (prevention, t_O,0_ = 0). Representative simulation results are shown in Figure 1B.

The study consisted of three parts. First, a stability analysis was performed to assess how astrocytic model parameters influence simulation stability. Second, the effects of these parameters on extracellular potassium dynamics and the resulting modulation efficacy were evaluated. Finally, the influence of the stimulation protocol and opsin properties on the effectiveness of optogenetic modulation was investigated.

### Analysis

The effectiveness of the optogenetic modulation in reducing [K^+^]_e_ can be described using the extracellular K^+^ concentration at the end of illumination ([K^+^]_e_(t_O,end_)) and via the relative change in [K^+^]_e_ at t = t_O,end_ between the no stimulation and stimulation conditions (Δ_rel_[K^+^]_e_).

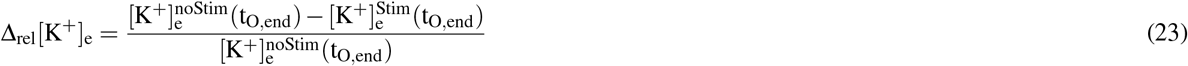

In the intervention paradigm the potassium clearance fraction (PCF) can also be used.

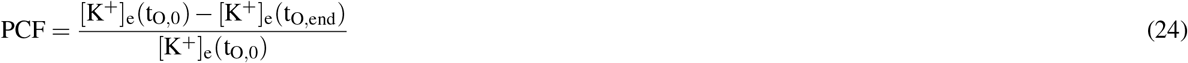

For the intervention setup, the potassium clearance time (PCT) is defined as the earliest time point between t_O,0_ and t_O,end_ at which [K^+^]_e_ falls within 1% of its minimum value, [K^+^]_e,min_, attained over this interval.

To determine the optimal duty cycle (dc)/pulse repetition period (prp) combination for the intervention stimulation paradigm, we identify the minimal duty cycle required to achieve near-maximal Δ_rel_[K^+^]_e_. Because this optimum depends on opsin kinetics, where small *τ*_on_ and large *τ*_off_ values are expected to yield the largest photocurrents, we separately evaluate the constraints imposed by opening and closing dynamics. For each duty cycle dc, we determine the maximal *τ*_on_ and the minimal *τ*_off_ for which Δ_rel_[K^+^]_e_ remains at at least 95% of its maximum. This defines two quantities: the *τ*_on_-limited minimum duty cycle, 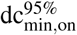, and the *τ*_off_-limited minimum duty cycle, 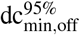, each representing the minimal duty cycle required to maintain near-maximal performance under the respective constraint.

To quantify the relationship between *τ* and 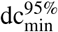, a linear regression model was fitted in log-log space using least-squares estimation^32^. Specifically, 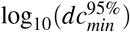 was modelled as a linear function of log_10_ (*τ*), yielding estimates of the regression coefficients. The fitted model was subsequently used to predict 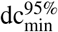 for a given *τ*_0_, after which predictions were transformed back to linear space by exponentiation. For each prediction, a 95% prediction interval (PI) was computed. The prediction interval was subsequently transformed back to linear space, resulting in asymmetric confidence bounds for 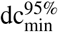. The critical duty cycle for a given prp was then defined as

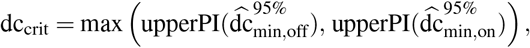

corresponding to a conservative estimate of the minimal duty cycle that simultaneously satisfies both *τ*_on_ and *τ*_off_ constraints.

## Results

### Astrocyte robustness

The nearly non-existent 100% percentile spread in Figure 2A indicates that neither the value of [K^+^]_e_ at t_O,0_ nor the effectiveness of astrocytic modulation strongly depends on the astrocyte’s volume Ω_A_ or surface to volume ratio *α*_A_ when *α*_ECS,A_ is held constant. This independence persists across all tested values of gap junction strength *σ*, maximal Na^+^*/*K^+^ ATPase pump current *ρ*, and KIR conductance g_KIR_, whereas *α*_ECS,A_ itself exerts a clear and significant influence. Astrocytic buffering is substantially stronger at higher *α*_ECS,A_ values, as reflected in both lower initial [K^+^]_e_(t_O,0_) and lower post-intervention [K^+^]_e_(t_O,end_) levels. The astrocytic parameters *σ, ρ*, and g_KIR_ also affect [K^+^]_e_ accumulation. Increasing these parameters consistently reduces [K^+^]_e_, although their relative impact depends on *α*_ECS,A_. For larger *α*_ECS,A_ values, the influence of *σ* and g_KIR_ becomes negligible compared to that of *ρ*. At the default *α*_ECS,A_ = 0.4, all parameters have a clear effect on both the initial and post-stimulation [K^+^]_e_.

**Figure 2.**
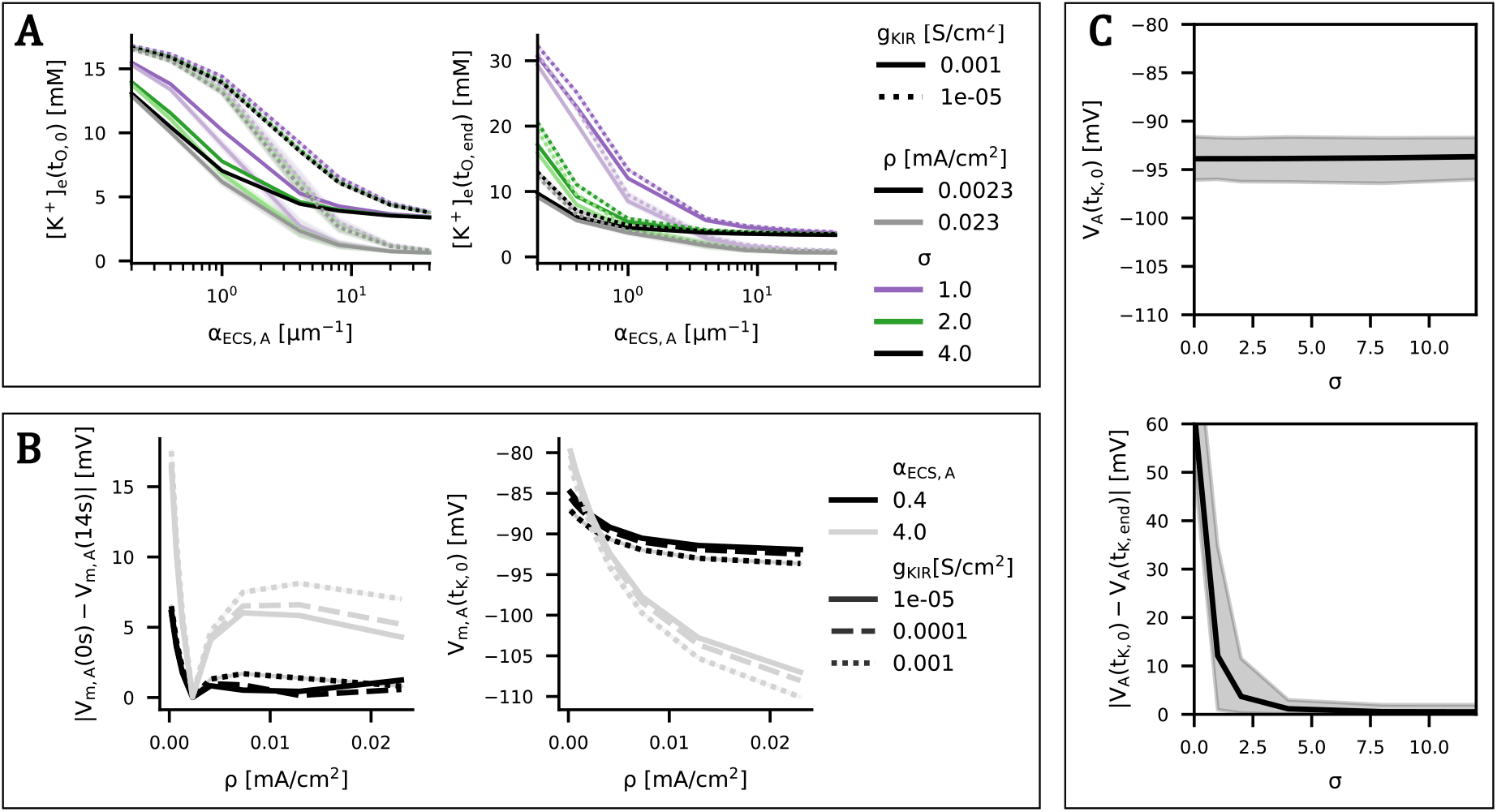
Impact of astrocyte model parameters on simulation stability using the Ksteady injection. **A**. Effect of astrocyte geometry on model output. The shaded region represents the full range (100% percentile) obtained by varying *α*_A_ (0.5–25 *µ*m^−1^) and Ω_A_ (2000–8000 *µ*m^3^). [K^+^]_e_(t_O,end_) shown for WiChR. **B**. Influence of gap-junction coupling strength *σ* on membrane stability. The shaded region represents the full range obtained by varying *ρ* (0.0023 - 0.023 mA*/*cm^2^), g_KIR_ (1e-5 – 1e-3 S*/*cm^2^), and K_inj_ (2.5 – 4 mM). No illumination. **C**. Influence of *α*_ECS,A_, *ρ*, and g_KIR_ on astrocytic membrane potential (V_m,A_) stability and steady-state value. No illumination and no K^+^ injection.

The default value of *ρ* was determined as the point at which the astrocytic membrane potential V_m,A_ remains stable when using P_K_ and P_Na_ values from Huguet et al. (2016) and Terman and Zhou (2019). Figure 2B shows that this value is independent of g_KIR_ and *α*_ECS,A_ and lies around 0.0025 mA*/*cm^2^. A more detailed analysis reveals that this value is independent of P_K_ but depends on P_Na_. Specifically, a linear relationship is observed between P_Na_ and the corresponding *ρ* required for V_m,A_ stability. The resulting linear regression (R^2^ = 0.9998) was used to determine the default value (*ρ* = 15 *×* 10^3^ P_Na_ + 10^−4^ = 0.0023mA*/*cm^2^).

When *α*_ECS,A_ = 0.4 and *ρ* values are above 0.0023mA*/*cm^2^, membrane potential deviations remain limited (within ~2.5 mV), and V_m,A_(t_K,0_) is largely independent of *ρ*, regardless of g_KIR_. In contrast for larger *α*_ECS,A_, V_m,A_ becomes more unstable (Fig 2B, left) and is more hyperpolarized (Fig 2B, right).

Figure 2C shows that V_m,A_ is unaffected by *σ* prior to K^+^ injection (V_m,A_(t_K,0_) stable). After injection, however, stronger gap junction coupling is required to maintain the isopotentiality necessary for inward K^+^ currents. Only *σ* ≥ 5 limits the increase in V_m,A_ to less than 5 mV, despite changes in E_K_.

### Impact of astrocytic parameter on potassium dynamics

Activation of WiChR in astrocyte A_0_ rapidly lowers [K^+^]_e_ for two different *ρ* values (fig 1B, a and b). In contrast, for ChR2^*^-based modulation, the decrease in [K^+^]_e_ is significantly less pronounced (1 mM decrease vs 3 mM in WiChR) and is even absent at low *ρ*, resulting only in a cessation of further potassium accumulation rather than an actual reduction (0.03 mM decrease vs 2.9 mM in WiChR). In addition, the effect of the ChR2^*^ photocurrent on the V_m,A_ is more pronounced than that of WiChR. Interestingly, optogenetic modulation decreases both intracellular and extracellular [K^+^] for both opsins. The intracellular reduction is likely driven by gap junction currents that redistribute K^+^ to the syncytium when V_m,A_ and/or [Na^+^]_i_ increase. Although [Na^+^]_i_ rises during both WiChR- and ChR2^*^-based modulation, the resulting changes in astrocytic ionic concentrations are considerably larger for ChR2^*^.

These observations are further supported by Figure 3A. For ChR2^*^, both Δ_rel_[K^+^]_e_ and potassium clearance fraction (PCF) values are substantially lower across all parameter ranges. When *σ* is low, Δ_rel_[K^+^]_e_ becomes negative, indicating an exacerbation rather than a reduction of potassium accumulation, while PCF values below zero indicate a failure to reduce [K^+^]_e_. The higher prevalence of positive values for Δ_rel_[K^+^]_e_ than for PCF suggests that, although ChR2^*^ can limit the rate of potassium accumulation, it is less effective at actively reducing already elevated [K^+^]_e_ levels.

**Figure 3.**
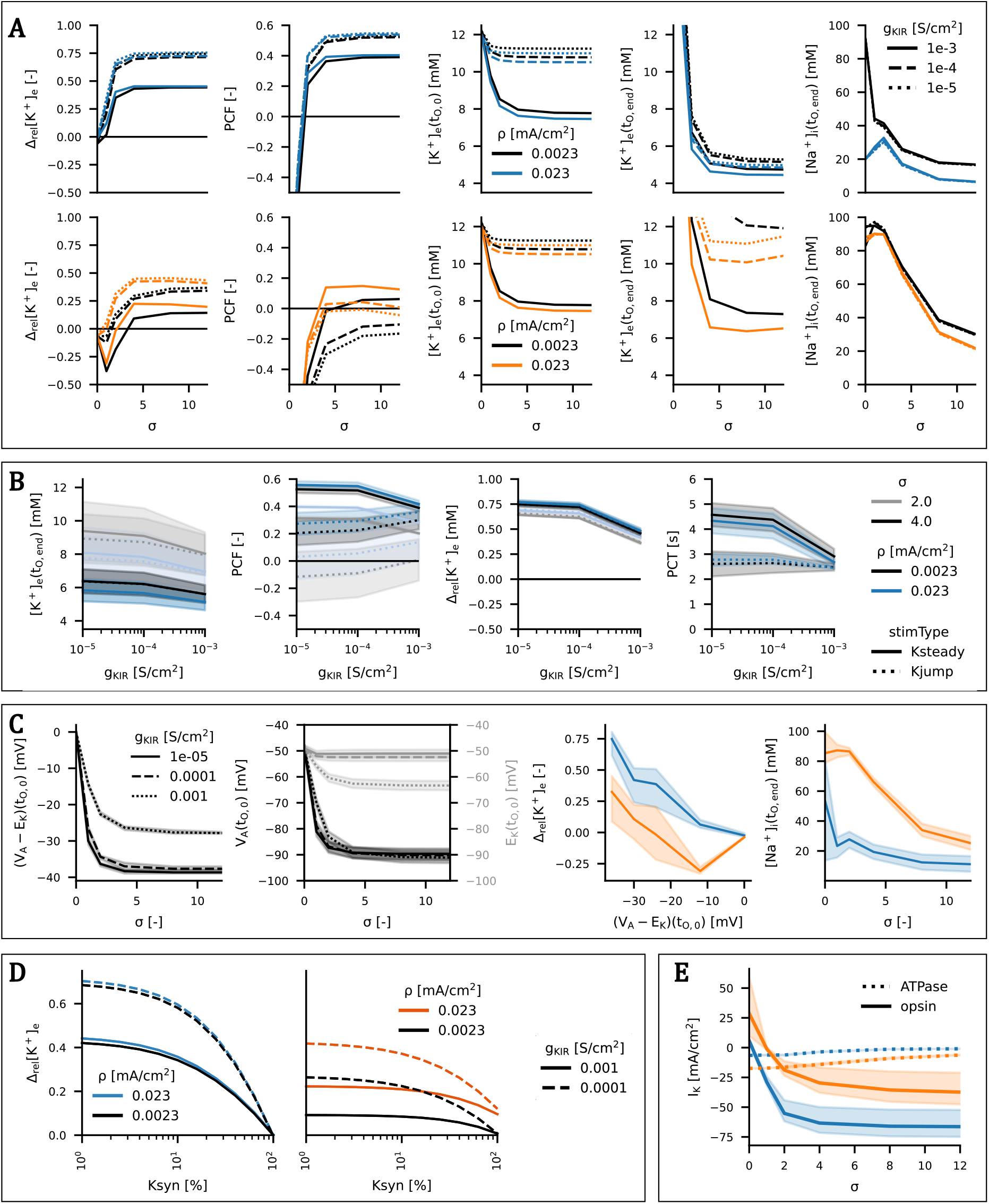
Results of astrocytic parameter analysis using WiChR 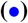 and ChR2^*^ 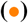 opsins. Parameter values: g_KIR_ = 1e-3/1e-4/1e-5 S*/*cm^2^, *ρ* = 0.0023/0.023 mA*/*cm^2^, *σ* = 0/1/2/4/8/12, and K_inj_ = 2.5/4 mM. All other parameters are set to their default values unless stated otherwise. **A**. Overview of the impact of astrocytic modulation across varying parameter combinations (K_inj_ = 2.5 mM, Ksteady). **B**. Detailed influence of g_KIR_ on WiChR-mediated modulation. The shaded region represents the full range (100% percentile) obtained by varying K_inj_. **C**. Influence of gap-junction coupling strength *σ* (K_inj_ = 2.5 mM, Ksteady) on isopotentiality, V_m,A_ E_K_ driving force and potassium clearance. The shaded region represents the full range obtained by varying all other parameters. **D**. Effect of increased syncytial potassium levels 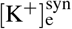 on optogenetic modulation. Δ_rel_[K^+^]_e_ is shown as a function of K_syn_, defined as the fraction of K_inj_ applied to A^syn^ (K_inj_ = 2.5 mM), for varying *ρ* and g_KIR_, with *σ* = 4, Ksteady. **E**. Partial potassium current contributions during illumination summed over time for varying *σ* (K_inj_ = 4 mM, *ρ ρ* = 0.023 mA*/*cm^2^). The shaded region represents the full range obtained by varying all other parameters.

The gap junction strength *σ* strongly influences astrocyte mediated potassium dynamics, both in the presence and absence of illumination (Figure 3A). For WiChR, increasing *σ* reduced extracellular potassium concentrations at both illumination onset ([K^+^]e(tO, 0)) and termination ([K^+^]e(tO, end)), resulting in improved performance according to the Δ_rel_[K^+^]_e_ and PCF metrics. This indicates that stronger coupling within the astrocytic syncytium enhances both natural and optogenetically mediated potassium buffering. A plateau in performance is observed for *σ ≥* 4 for all WiChR metrics. In contrast, ChR2^*^ performance peaks around *σ* = 4 for certain astrocyte configurations and declines at higher values. Figures 3C and E clarify this behaviour: increasing *σ* enhances syncytial isopotentiality, stabilizing V_m,A_ at illumination onset and increasing the driving force (V_m,A_ − E_K_) for potassium currents. For both opsins, a larger driving force leads to a stronger Δ_rel_[K^+^]_e_ through increased inward potassium current. However, ChR2^*^ additionally significantly elevates intracellular Na^+^ (Figure 3C), which enhances the Na^+^*/*K^+^-ATPase pump current. As shown in Figure 3E, the photocurrent-to-pump-current contribution to potassium regulation is much larger for WiChR than for ChR2^*^, making ChR2^*^ more dependent on pump activity. At high *σ*, the increase in [Na^+^]_i_ is reduced, leading to diminished pump activation and decreased ChR2^*^ efficacy. The K^+^ current contributions also show that ChR2^*^ can generate an inward K^+^ current when *σ* is high. Furthermore, these contributions help explain why ChR2^*^ modulation can increase [K^+^]_e_ (Δ_rel_[K^+^]_e_ *<* 0): under conditions of low coupling, depolarizing mixed K^+^ and Na^+^ ChR2^*^ currents can raise V_m,A_ above E_K_, thereby reversing the direction of the potassium current and promoting outward K^+^ flux.

The parameter *ρ* has minimal influence on WiChR-mediated modulation but significantly affects ChR2^*^, consistent with its reliance on pump-mediated potassium removal (Figure 3A).

The impact of g_KIR_ is illustrated in Figures 3A and B. Higher g_KIR_ values lead to lower [K^+^]_e_(t_O,end_) but also reduce PCF and Δ_rel_[K^+^]_e_ in WiChR, an effect attributable to differences in [K^+^]_e_(t_O,0_) that confound the comparison between g_KIR_ values. This confounding effect is avoided in the Kjump protocol, where potassium injection starts only at t_O,0_ and initiates with an instantaneous increase of [K^+^]_e_ to 8mM. Under these conditions, PCF increases with increasing g_KIR_. This suggests that higher g_KIR_ does not make the modulation less effective as the decrease is in PCF and Δ_rel_[K^+^]_e_ would suggest but indicates that there is a lower bound in achievable [K^+^]_e_ under stimulation that requires a larger reduction when g_KIR_ is low and causing high initial concentrations. Lower g_KIR_ also slows potassium clearance under Ksteady (decreased PCT). This can be attributed to the higher [K^+^]_e_ at illumination start as this dependency is absent under Kjump conditions, where PCT becomes independent of g_KIR_.

Across variations in K_inj_ and initial [K^+^]_e_, Figure 3B further shows that Δ_rel_[K^+^]_e_ remains robust for both injection paradigms, indicating that this metric is suitable for consistent comparison across astrocytic and stimulation parameters.

Overall, improved astrocytic potassium-buffering capacity, characterised by higher g_KIR_ and *ρ*, enables lower potassium levels at then end of optogenetic modulation. However, when large potassium elevations occur in well-functioning astrocytes, they are more difficult to counteract due to the higher potassium load (Figure 3A). Increasing *σ* consistently enhances buffering and improves WiChR-mediated modulation, largely independently of g_KIR_ and *ρ*, as reflected by the near-parallel behaviour across parameter combinations. In contrast, excessively high *σ* can reduce ChR2^*^ effectiveness due to diminished [Na^+^]_i_ accumulation and consequently weaker ATPase pump activation.

Figure 3D illustrates the effect of widespread potassium elevation within the syncytium. The ability of WiChR to reduce [K^+^]_e_ diminishes as the ratio 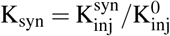 increases, and disappears when K_syn_ = 1. This occurs because 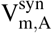 approaches 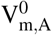, eliminating the stabilising effect of syncytial coupling and reducing the driving force for the potassium photocurrent current as V_m,A_ follows E_K_. In contrast, for ChR2^*^, modulation does not vanish when the ATPase pump is sufficiently strong. The higher E_rev_ associated with lower P_K*/*Na_ maintains a driving force (V_m,A_ − E_rev_), sustaining inward sodium currents. The resulting increase in [Na^+^]_i_ continues to enhance pump activity, allowing residual modulation under these conditions.

In conclusion, WiChR provides more consistent and effective potassium regulation than ChR2^*^, except in conditions where syncytial coupling is compromised and ATPase activity is sufficiently strong to support ChR2^*^-mediated modulation.

### Impact of stimulation and opsin parameters on potassium dynamics

Figure 4A illustrates the impact of the optogenetic stimulation paradigm on potassium modulation. The results show that light intensity I is a key determinant of modulation outcome. For I *≥* 0.1mW*/*mm^2^, a plateau is reached beyond which Δ_rel_[K^+^]_e_ no longer improves. When dc = 0.001, Δ_rel_[K^+^]_e_ decreases substantially, but increases in duty cycle beyond 0.01 have only a limited effect. However, the influence of dc becomes more pronounced at higher prp values, where lower duty cycles lead to smaller reductions in Δ_rel_[K^+^]_e_. Lower prp values yield better performance for the same energy usage, as the slow closing kinetics of the opsin allow more frequent pulses to generate a larger cumulative photocurrent. Figure 4B confirms this relationship, showing that larger total photocurrents correspond to higher effectiveness (larger Δ_rel_[K^+^]_e_), while larger prp values result in reduced total photocurrent. The difference between the intervention (t_O,0_ = 7.5s) and prevention (t_O,0_ = 0s) paradigms remains below Δ_rel_[K^+^]_e_ = 0.005 for both WiChR and ChR2^*^ across all parameter combinations, indicating a lower bound for extracellular potassium regulation that is largely independent of stimulation timing.

**Figure 4.**
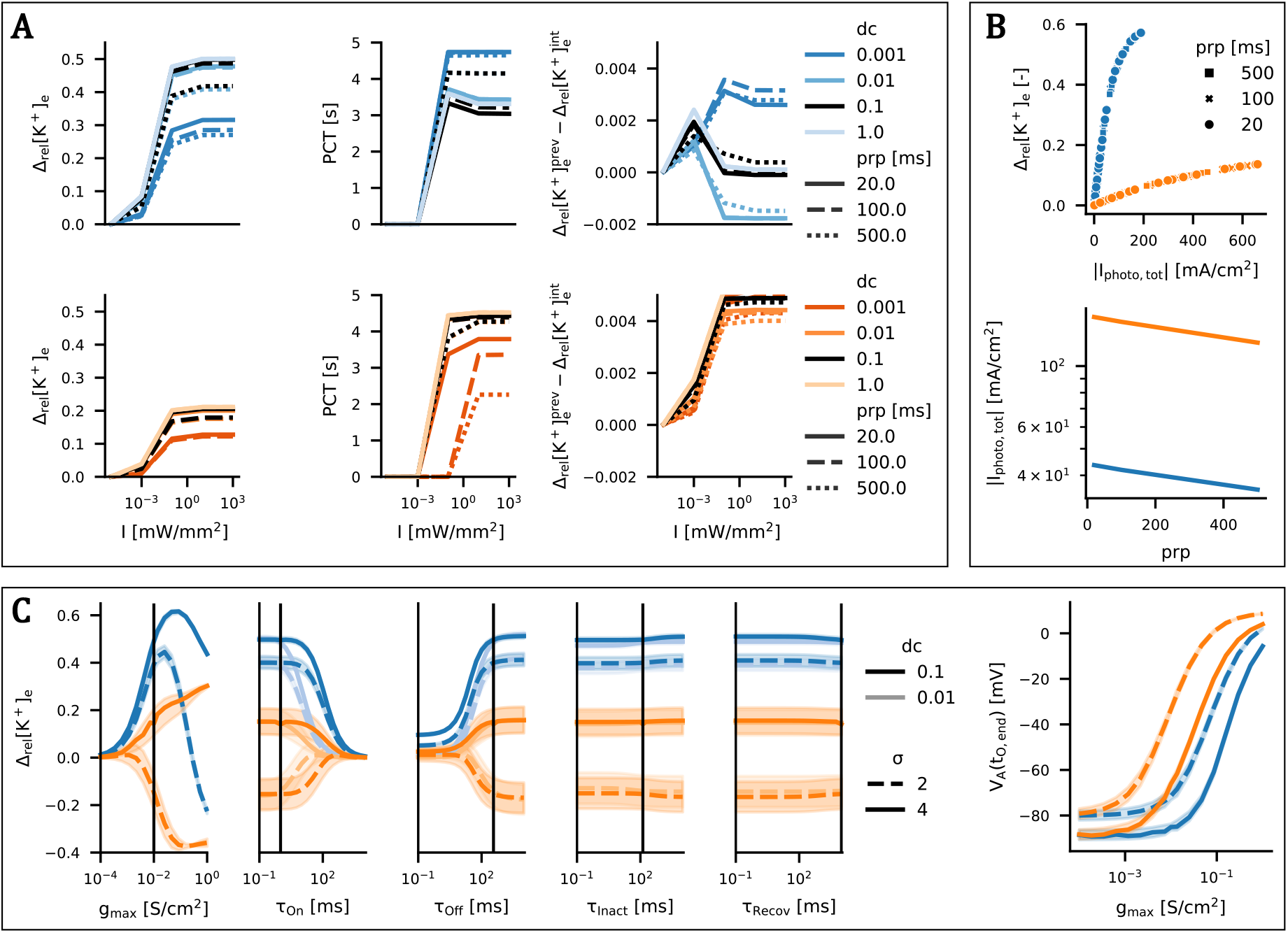
Impact of opsin and stimulation parameters on potassium dynamics using WiChR 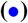 and ChR2^*^ 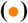. **A**. Influence of stimulation parameters for WiChR (*ρ* = 0.0023 mA*/*cm^2^, 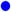) and ChR2^*^ (*ρ* = 0.023 mA*/*cm^2^, 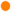. Astrocytic parameters are set to default values. **B**. Influence of prp on modulation outcome and total photocurrent. **C**. Effect of opsin parameters on Δ_rel_[K^+^]_e_ for varying dc and *σ*. The shaded region represents the full range obtained for *ρ* = 0.0023 and 0.023 mA*/*cm^2^. All other parameters are set to their default values.

Intrinsic opsin properties also influence modulation effectiveness. Figure 4C isolates the impact of individual parameters. While larger g_max_ values might be expected to improve performance due to the higher generated photocurrents, WiChR exhibits an optimal g_max_, beyond which effectiveness decreases. This reduction is likely due to increased depolarization of the astrocytic membrane (Figure 4C, right), which raises V_m,A_ toward E_K_ and thereby reduces the driving force for potassium currents following an initial increase. Because lower *σ* values correspond to higher V_m,A_, the optimal g_max_ shifts to lower values under weaker coupling. In contrast, ChR2^*^ does not exhibit such an optimum, likely because increased Na^+^-driven pump activity continues to contribute to potassium buffering even when the direct photocurrent diminishes. This effect is not observed when gap-junction coupling is reduced. Under these conditions, increasing ChR2^*^ g_max_ leads to greater potassium accumulation rather than enhanced clearance, likely because the resulting higher depolarizing current raises V_m,A_ above E_K_, thereby reversing the direction of the potassium current and promoting outward potassium flux.

The inactivation and recovery time constants have minimal impact on modulation outcome, whereas the opening and closing time constants play a more substantial role. When *τ*_on_ is large, channels do not fully open during short illumination pulses, reducing effectiveness. This effect is exacerbated at lower dc. Conversely, when *τ*_off_ is too small, rapid channel closing shortens the photocurrent tail, reducing total current and requiring compensation via increased dc.

These dependencies define an optimal effective illumination duration (determined by dc *·* prp), which depends on the opsin kinetics. To quantify this, 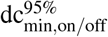 was computed as a function of *τ*_on_ and *τ*_off_ for different prp values (Figure 5A), representing the minimal dc required to achieve at least 95% of maximal Δ_rel_[K^+^]_e_. However, this metric can only be defined when the *τ*_on_-Δ_rel_[K^+^]_e_ curve follows the behaviour shown in Figure 4C. As shown in Figure 5A (right), this condition is not always satisfied: for sufficiently low dc *·* prp values, performance collapses to zero regardless of *τ*_on_. This introduces a hard lower limit dc_0_ for each prp (Figure 5B).

**Figure 5.**
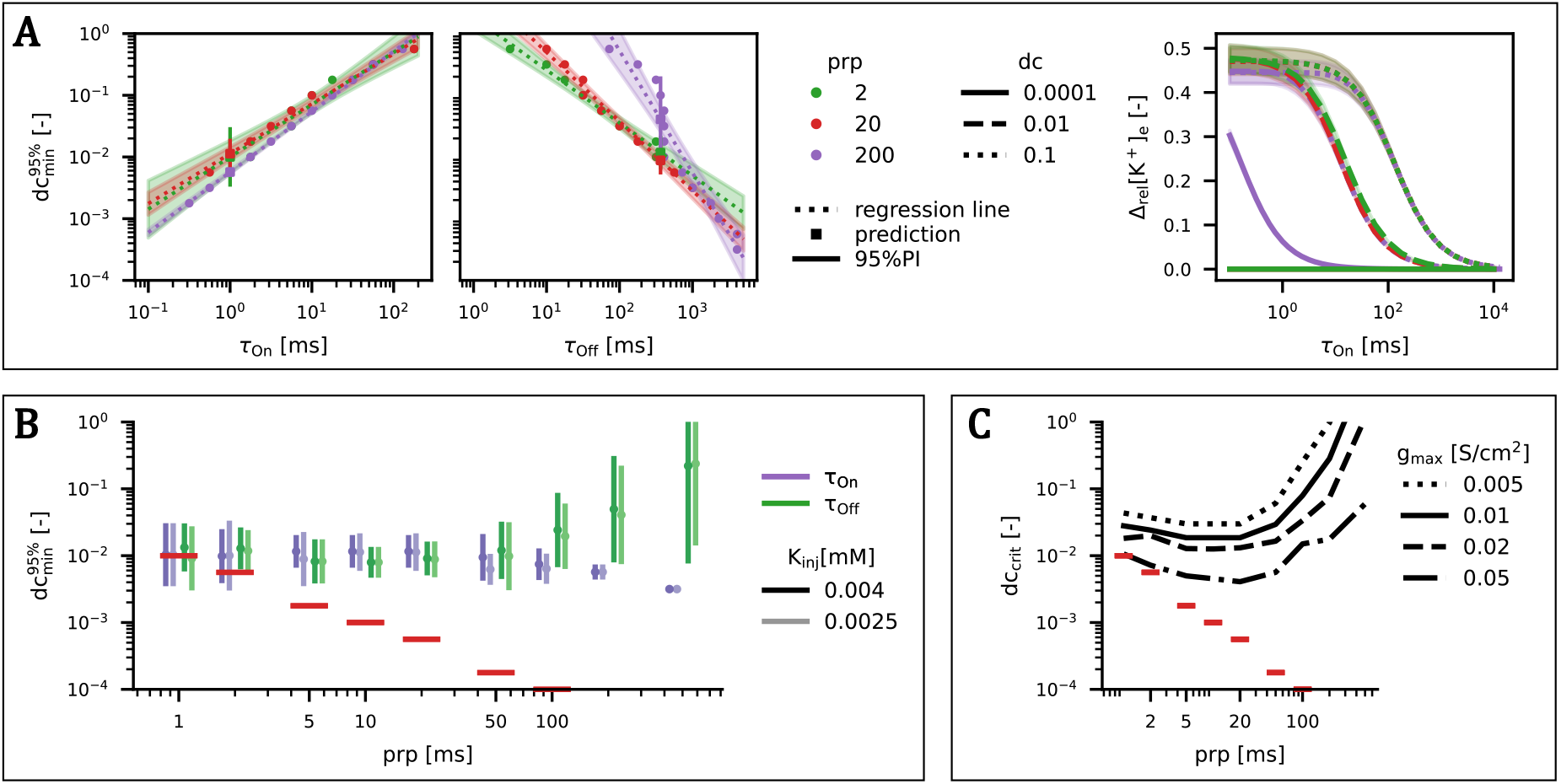
Optimization of stimulation parameters. **A**. 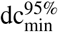 as a function of *τ*_on_ and *τ*_off_ for varying prp (left). Δ_rel_[K^+^]_e_ as a function of *τ*_on_ for varying prp (right). (I = 10^3^ mW*/* mm^2^, g_max_ = 10^−2^ S*/*cm^2^) **B**. PI95% of 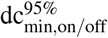 as a function of prp for *τ*_on_ = 1 ms and *τ*_off_ = 365 ms. The red lines indicates dc_0_ = hard lower limit for each prp regardless of *τ*_on_. **C**. dc_crit_ as a function of prp for varying g_max_ (*τ*_on_ = 1 ms, *τ*_off_ = 365 ms). The red markers indicate dc_0_.

For parameter combinations where 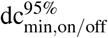 is defined, a log-log linear relationship is observed, enabling prediction of 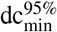 across *τ*_on_ and *τ*_off_ values. Applying this model to WiChR’s kinetic parameters yields the prediction intervals shown in Figure 5B, which appear independent of K_inj_ and can be used to define a critical duty cycle dc_crit_. As shown in Figure 5C, dc_crit_ depends on both prp and g_max_. Notably, the optimal prp is approximately 20 ms, independent of g_max_, while higher g_max_ values correspond to lower required dc_crit_.

## Discussion

### Model assumptions and parameterisation

The present study is based on a conductance-based computational framework designed to isolate the dominant ionic mechanisms underlying astrocyte mediated potassium dynamics during optogenetic modulation. Within this framework, WiChR and ChR2 are modelled as cation conducting channels with different permeability ratios for sodium and potassium, with WiChR exhibiting approximately 60 times higher potassium selectivity. As a result, ChR2 influences potassium dynamics both directly and indirectly via increased intracellular Na^+^ concentration and subsequent boosting of the Na^+^*/*K^+^-ATPase pump.

Although ChR2 is permeable to multiple ionic species, including H^+^ and Ca^2+^, these contributions were neglected. The effect of H^+^ is negligible under physiological conditions due to its low concentration and previous modeling studies indicate that optogenetic modulation of astrocytes is dominated by Na^+^ and K^+^ dynamics^17,28^. The model therefore focuses on these ions to enable systematic analysis of potassium regulation. Such simplifications are necessary to facilitate systematic exploration of the parameter space while retaining the key processes governing extracellular potassium dynamics. For this same reason, the parameters describing the opsin dynamics were chosen to be the same for both WiChR and ChR2^*^.

The astrocytic syncytium, which in reality consists of hundreds of interconnected astrocytes, is represented by a single compartment. Consequently, the coupling parameter *σ* captures both gap-junction strength and the aggregate effect of multiple coupled astrocytes. Because Ma et al. (2016) estimated a value of approximately 1 for coupling between individual astrocytes, this estimate cannot be directly applied to the present model. Therefore, *σ* is treated as an uncertain parameter and is allowed to exceed 1^17^. The default value was determined using the experimental observation that syncytial isopotentiality attenuates membrane depolarization induced by local increases in extracellular potassium. Specifically, syncytial coupling reduces the astrocytic membrane potential change induced by a local elevation of [K^+^]_e_ of 23 mM from 43 mV to 5 mV^17^. In the model, this behaviour is reproduced only for *σ ≥* 4, for which the resulting change in V_m,A_ remains below 5 mV when seizure-associated levels of potassium are reached ([K^+^]_e_ *>* 8 mM) (Figure 2C).

The surface-to-volume fractions *α*_ECS,A_ and *α*_A_ determine the extent to which astrocytic membrane currents influence extra- and intracellular ion concentrations, respectively. As long as these ratios are preserved, absolute compartment volumes are not expected to affect the dynamics. Astrocytes occupy approximately 5-10% of brain tissue, while the extracellular space comprises roughly 6-20%, implying a broad range of possible values for 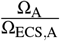 (0.25-1.67)^33,34^. Experimental estimates place the astrocytic surface-to-volume ratio *α*_A_ between 15 and 25 *µ*m^−1^, whereas computational studies have used values as low as 0.8^17,22,28,30^. These estimates yield 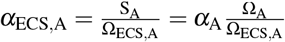 values ranging from approximately 4 to 40 *µ*m^−1^, or as low as 0.2 *µ*m^−1^ when computational estimates are considered^17^. Astrocytes have been reported to clear extracellular potassium at rates of up to 2 mM/s^16^. Guided by this estimate, a default value of *α*_ECS,A_ = 0.4 *µ*m^−1^ was selected as a reference condition. This value corresponds to the threshold at which an astrocyte with default parameters (from Terman and Zhou (2019) and Huguet et al. (2016)) is unable to prevent [K^+^]_e_ from reaching seizure-associated levels (8 mM) by t_O,0_ under a potassium injection rate of K_inj_ = 2.5 mM/s. This reference therefore provides a physiologically motivated baseline within the broader parameter sweep. Further parameter analysis showed that the model is primarily sensitive to *α*_ECS,A_ and the absolute value of *α*_A_ has little effect provided the relevant astrocyte surface to ecs volume ratio (*α*_ECS,A_) is preserved (Figure 2A).

### Mechanistic comparison of WiChR and ChR2-based modulation

The comparison between WiChR and ChR2^*^ reveals fundamentally distinct mechanisms of astrocyte mediated potassium regulation. In agreement with Zhao et al. (2022)^10^, ChR2-based modulation relies strongly on Na^+^*/*K^+^-ATPase activity, as reflected by the sensitivity to *ρ* observed both *in vivo* and *in silico*. In contrast, little dependence on g_KIR_ is observed, consistent with previous experimental and computational findings showing that Kir4.1 blockade increases seizure occurrence but does not abolish the modulatory effect of ChR2^10,35^. Furthermore, inhibition of gap junctions reverses anti-seizure effects *in vivo*, while the model highlights the requirement for sufficiently strong coupling (high *σ*).

A key insight of the present work is that WiChR provides more consistent and robust potassium regulation than ChR2^*^. By directly mediating K^+^ flux, WiChR reduces the need for large intracellular sodium increases and working Na^+^*/*K^+^-ATPase pumps, which may be impaired under pathological conditions^36,37^. Although ChR2^*^ relies on potassium removal through sodium-driven pump activation, potassium current decomposition indicates that its partial potassium permeability can, under certain conditions, contribute directly to potassium clearance, occasionally exceeding the contribution of pump activation and revealing an additional mechanism that was not fully considered in previous work^10^. Under conditions of compromised gap junction coupling, ChR2^*^ can even exacerbate potassium accumulation due to depolarisation-induced outward potassium currents, whereas WiChR’s behaviour is more robust and predictable.

The model identifies gap-junction coupling (*σ*) as a critical determinant of astrocyte-targeted modulation, as it enables both effective potassium redistribution and the maintenance of syncytial isopotentiality which sustains the driving force for potassium uptake. Notably, even when gap junction coupling is impaired, WiChR does not induce a net increase in [K^+^]_e_. This is because the WiChR current consists almost exclusively of K^+^, causing V_m,A_, E_K_ and E_rev_ to almost overlap, and thereby reducing the driving force toward zero. In contrast, for ChR2, the reversal potential E_rev_ is higher than E_K_, such that V_m,A_ *>* E_K_ is maintained, resulting in an outward potassium photocurrent that can promote potassium accumulation. Furthermore, for ChR2-based modulation, an optimal range of *σ* emerges. This highlights the dual role of the syncytium in maintaining electrochemical gradients while enabling spatial buffering of potassium. As astrocytic dysfunction is known to occur in epilepsy, ranging from gap junction uncoupling to altered connexin expression, the effectiveness of astrocyte targeted modulation may depend strongly on disease or patient specific alterations in gap junction coupling^38–40^.

The model results also indicate that WiChR-based modulation critically depends on ion homeostasis within the syncytium: when extracellular potassium is elevated throughout the entire syncytium, the modulation effect is lost. Under these conditions, V_m,A_ approaches E_K_, eliminating the driving force for inward potassium photocurrents. In contrast, ChR2-based modulation can remain effective, provided that Na^+^*/*K^+^-ATPase activity is sufficiently strong, as sodium-driven pump currents continue to support potassium clearance.

Finally, both optogenetic strategies can induce astrocytic depolarisation. This effect is more pronounced for ChR2^*^ (Figure 4C, right) and may influence voltage-dependent pathways including glutamate clearance and intracellular calcium signalling^18^. Astrocytic depolarisation has been shown to reduce glutamate uptake and increase neuronal excitability, while also influencing Ca^2+^-dependent gliotransmission and cerebral blood flow^37,41–44^. More detailed astrocyte models incorporating intracellular calcium dynamics and neuron-astrocyte signalling have previously been developed to investigate these interactions in greater detail^35,45^. The present work, however, was designed to focus specifically on potassium-mediated mechanisms of seizure modulation, and therefore did not include explicit calcium dynamics. This simplification is supported by the findings of Maboodi et al., who reported that seizure suppression by astrocytic ChR2 activation remained largely preserved despite impaired astrocytic calcium signalling, suggesting that ion-homeostatic mechanisms rather than calcium-dependent pathways constitute the dominant anti-seizure effect^35^. Nevertheless, the model cannot assess potential contributions of calcium-dependent astrocytic signalling to seizure dynamics and recovery.

### Translational implications and stimulation parameter optimisation

The computational findings suggest that direct modulation of extracellular potassium via facilitation of K^+^ photocurrents in astrocytes could represent a viable strategy to reduce pathological increases of extracellular potassium and prevent or stop epileptic seizures. From a translational perspective, optimisation of stimulation parameters is particularly important, as low-power operation is desirable to minimise tissue heating and energy consumption in implantable systems.

The model indicates that optimal stimulation parameters depend strongly on intrinsic opsin properties, such as opening and closing kinetics, as well as on g_max_. In particular, the prediction that pulse repetition period (prp) values around 20 ms yield near-optimal performance provides a useful starting point for in vivo protocol design. This optimum arises because intermediate prp values balance two competing effects: at high prp, the photocurrent tail is not efficiently utilised between pulses, while at low prp, channels do not have sufficient time to fully open. As a result, optimal performance is achieved for prp values in the mid-range.

The optimal duty cycle (dc) is more difficult to determine, as it depends on g_max_, which is challenging to estimate experimentally. This parameter depends on both opsin expression level and single-channel conductance, which are rarely quantified and reported estimations for ChR2 vary over several orders of magnitude^21,25^. However, in vitro experiments measuring the amplitude and temporal dynamics of potassium changes during illumination could provide constraints on model parameters and enable more accurate estimation of g_max_.

### Conclusion and future perspectives

In this study, we systematically investigated astrocyte-mediated optogenetic modulation of extracellular potassium using potassium-selective (WiChR) and cation-conducting (ChR2-like) opsins within a unified computational framework. Our results demonstrate that direct modulation of K^+^ dynamics via WiChR provides a more robust and consistent mechanism for reducing [K^+^]_e_ than indirect, pump-dependent strategies based on ChR2^*^. This advantage arises from the ability of WiChR to directly enhance astrocytic potassium buffering, largely independent of Na^+^*/*K^+^-ATPase activity and with reduced sensitivity to variations in astrocytic parameters. Furthermore, modulation efficacy emerges from the interplay between opsin properties and stimulation parameters, with an optimal pulse repetition period of approximately 20 ms observed largely independent of g_max_. This finding provides a practical and robust guideline for stimulation design. These findings provide a foundation for the rational design of astrocyte-targeted optogenetic interventions aimed at seizure suppression.

Importantly, our results also indicate that modulation efficacy depends on the integrity and homeostasis of the astrocytic syncytium, highlighting the need to better characterise astrocyte network properties and their alterations under pathological conditions. Future work should focus on experimental validation, improved parameter estimation, and the incorporation of additional physiological processes. In particular, measurements of extracellular potassium dynamics during optogenetic stimulation would enable validation and calibration of the model predictions. While the present study focused on potassium-mediated mechanisms, future models could incorporate calcium signalling and gliotransmission pathways to investigate potential secondary consequences of astrocytic depolarisation that were beyond the scope of the current work.

Beyond epilepsy, the ability to modulate extracellular potassium dynamics may have broader applications in neurological disorders associated with potassium dysregulation, including spreading depolarisation, migraine, chronic pain, and possibly neurodegenerative diseases such as Alzheimer’s disease^28,46–49^. The integration of mechanistic modelling with experimental validation offers a promising framework for the development of targeted neuromodulation strategies across these diverse pathological contexts.

## Author contributions

LW: Conceptualization, Formal Analysis, Methodology, Software, Visualization, Writing – Original Draft

TT: Conceptualization, Methodology, Writing – Review & Editing.

WJ: Funding Acquisition, Writing – Review & Editing.

ET: Conceptualization, Methodology, Funding Acquisition, Writing – Review & Editing.

RR: Conceptualization, Methodology, Funding Acquisition, Writing – Review & Editing.

## Acknowledgements

This research was supported by the Ghent University Special Research Fund BOF-GOA project SOFTRESET (grant number 01G02722).

